# Natural isotope correction improves analysis of protein modification dynamics

**DOI:** 10.1101/2020.10.31.361725

**Authors:** Jörn Dietze, Alienke van Pijkeren, Anna-Sophia Egger, Mathias Ziegler, Marcel Kwiatkowski, Ines Heiland

## Abstract

Stable isotope labelling in combination with high resolution mass spectrometry approaches are increasingly used to analyse both metabolite and protein modification dynamics. To enable correct estimation of the resulting dynamics it is critical to correct the measured values for naturally occurring stable isotopes, a process commonly called isotopologue correction or deconvolution. While the importance of isotopologue correction is well recognized in metabolomics, it has received far less attention in proteomics approaches. Although several tools exist that enable isotopologue correction of mass spectrometry data, the majority is tailored for the analysis of low molecular weight metabolites. We here present PICor which has been developed for isotopologue correction of complex isotope labelling experiments in proteomics or metabolomics and demonstrate the importance of appropriate correction for accurate determination of protein modifications dynamics, using histone acetylation as an example.

## Introduction

Protein modifications such as phosphorylation, methylation or acetylation have an important regulatory function in biology and are the basis of environmental adaptation and phenotypic variations. While it is well known that protein phosphorylations are highly dynamic and result from a complex interplay between protein kinases and phosphatases, the dynamics of other protein modifications such as methylation and acetylation have been less well investigated. The availability of stable isotope labelled compounds and high resolution mass spectrometry has, however, more recently paved the way to analyse the dynamics of protein methylation, acetylation, phosphorylation and glycosylation^1–5^.

As basically all protein modifications are effectuated by small metabolic compounds, one can use partially or fully labelled metabolic precursors to monitor changes in concentrations of both metabolic intermediates and protein modifications over time. A typical experimental workflow for stable isotope labelling experiments starts with the incubation of cells with labelled precursors. Metabolites and proteins are then extracted at various time points, separated by gas chromatography (GC) or liquid chromatography (LC) and analyzed by mass spectrometry (MS).

Molecules with the same composition of isotopes, so-called isotopomers, are chemically identical. Their intact precursor ions are isobaric and therefore indistinguishable by MS analysis. An additional fragmentation by tandem mass spectrometry (MS/MS or MS2) using e.g. collision induced dissociation (CID), allows to localize the isotopes based on their presence in the individual fragments. The obtained data need to be appropriately processed, in order to correctly calculate metabolite or protein modification dynamics^6^. A further complication in this analysis is the necessity to correct the obtained values for natural isotope abundance, called isotop(ologu)e correction or deconvolution. This is critical, because naturally occurring compounds already contain 1.1% of the stable heavy isotope ^13^C. Thus, integration of ^13^C-labelled precursors requires the correction for this naturally occurring isotope and potentially also for other stable natural isotopes such as deuterium, ^18^O, or ^15^N, depending on whether or not these isotopes can be resolved by the detection method used^7^. Especially the assessment of large metabolites as well as peptides and proteins could otherwise lead to considerable errors in the estimation of the label integration dynamics.

In recent years, a number of tools have been made available that enable isotopologue correction for metabolites. But very little attention has been paid to natural isotope correction in proteomics or for the analyses of protein modification turnover. The theoretical basis and application of available tools for metabolites varies. Correction can for example be done based on spectra from naturally occurring compounds. This approach is, however, not feasible for proteomics applications, as it would require to have all peptides of interest available as standards, let alone inclusion of protein modifications. Similarly unfeasible for proteomics-related measurements is the application of tools that require the compilation of a list containing all expected isotopologues as is the case for example for IsoCor^8^.

The approach we chose here corrects measured values based on the calculated theoretical isotope distribution. The latter can be described using probability matrices. Commonly, two different types of matrices are used for natural isotope correction^9–11^: In the so-called *classical* approach, the correction matrix is based on the same spectrum or distribution for all measured isotopologues of the same atom composition. This is also the basis for correction methods based on measured spectra from naturally occurring substances. In molecules, however, that have all light isotopes replaced by their heavy counterparts, no heavier isotopologues can exist. These methods therefore lead to an estimation bias towards heavy isotopologues especially for large molecules with high label incorporation. We therefore used a so-called *skewed* matrix approach, that corrects each isotoplogue with its own set of theoretical calculated correction factors. This method is for example implemented in the R-based toolbox IsoCorrectoR^12^ and in AccuCor^7^.

For the identification and quantification of isobaric modified histone species, Yuan et al.^13^ developed an alternative method for isotope correction using a deconvolution approach based on MS/MS measurements. The corresponding toolbox EpiProfile has been primarily developed to analyse isobaric modified histone peptides irrespective of whether they arise from label integration or are caused by similar chemical composition. As this method allows to identify which part of the molecule is labelled with heavy isotopes it does enable some correction for natural isotope abundance. But, beside the disadvantage of requiring MS/MS measurements, it does not provide a full isotopologue correction as it ignores natural labelling of the peptide backbone. The resulting estimation error is increasing with the number of possible isotopologues and the number of atoms labelled.

We have therefore developed PICor (https://github.com/MolecularBioinformatics/PICor), a Python based isotopologue correction tool that is applicable for the analysis of both labelled metabolites as well as peptides and protein modifications. It enables the correction for incorporation of multiple different isotopes for example both ^13^C and ^15^N and uses the skewed matrix approach. Importantly, our approach is based on the theoretical distribution of isotopologues. Therefore, only the most abundant or important isotopologues need to be measured and thus incomplete datasets can be used. Using time course measurements of acetylated histone peptides we demonstrate that appropriate correction for natural isotope distribution considerably improves the accuracy of the analyses.

## Results and Discussion

We here demonstrate the effect of natural isotope correction on labelling experiments in proteomics and metabolomics by analysing MS data from ^13^C-glucose supplementation time courses. In these metabolic labelling experiments, naturally occuring ^12^C-glucose is replaced by ^13^C-glucose in cell culture media. In the cells the labelled glucose is, among others, converted into ^13^C-acetyl-Coenzyme A (acetyl-CoA). As acetyl-CoA is also the substrate for histone acetylations, natural occuring ^12^C-acetylation of lysines are replaced with ^13^C-labelled acetyl-groups causing a characteristic mass shift of the peptide in high resolution MS-spectra. The time dependent replacement of ^12^C-acetylation can be used to determine histone acetylation dynamics^1–3,6^.

All currently available tools for isotopologue correction have been developed for the correction of metabolite data but not all tools can correct for the incorporation of multiple different isotopes, like ^13^C and ^15^N. Those that can account for the latter are not suitable for the use in proteomics approaches as they either only allow for full or no resolution of different isotopologues (IsocorrectoR) or do require the full reporting of all possible isotopologues (IsoCor). For an overview of the properties and limitations of different available tools see Table 1. We have therefore developed a new Python based isotopologue correction tool PICor that enables the correction of both metobolomics and proteomics measurements. To show that the calculations of PICor are in agreement with other existing tools, we first analysed measurements from ^13^C labelled acetyl-CoA (Figure 1) as some of the currently available tools used for the method comparison can only be used for metabolites.

**Table 1:** Properties, and application areas of different tools for isotopologue correction. (Abbr.: Adj: high resolution mode with resolution dependent correction; P: proteomics; M: metabolomics.)

|  | PICor | IsoCorrectoR | AccuCoR | IsoCor |
| --- | --- | --- | --- | --- |
| Programming language | Python | R | R | Python |
| Focus | P & M | M | M | M |
| Multiple isotope labels | Yes | Yes | No | No |
| Resolution | Adj. | Low & High | Adj. | Low & Adj. |
| Correction matrix | Skewed | Skewed | Skewed | Classic |

**Figure 1:**
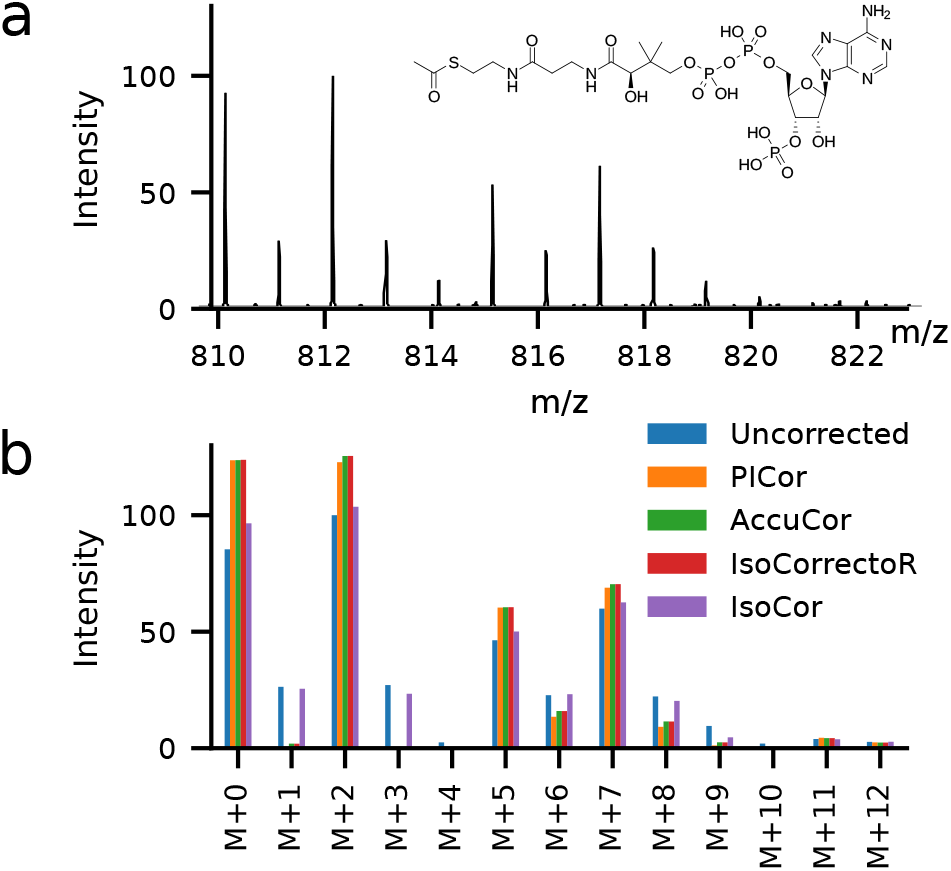
Metabolic labelling of acetyl-CoA with ^13^C-glucose after 6 h. a) Measured spectrum.b) Comparison of uncorrected peak heights from the spectrum shown in a) and values derived through correction with PICor, AccuCor, Iso-CorrectoR and IsoCor.

Acetyl-CoA is a relatively large metabolite, but the number of possible isotopologues is still much more limited than for peptides. As shown in Figure 1b the corrected spectra derived with PICor match those of other tools that use the skewed matrix approach, like IsoCorrectoR and AccuCor. The results do, however, clearly differ from the correction based on the classical matrix correction (e.g when using IsoCor).

When correcting for natural isotopes it is important to consider the mass resolution of the experiments^7^. Nowadays, it is common to use a measurement resolution that can separate ^13^C and ^15^N. But, resolving the mass difference between ^13^C and deuterium would require a resolution that is unfeasible for proteomics approaches in practice, due to required measurement time. PICor, therefore, enables isotopologue correction for all label combinations according to the mass resolution used in the experiments, making it especially relevant for labelling experiments using combinations of different isotopes. To make PICor applicable for proteomics approaches one only needs to provide a table with measurements for the isotopologues of interest and not a full list for all possible isotopologues as for example required for the analysis with IsoCor.

### Isotopologue correction is important to correctly estimate histone acetylation dynamics

Histone acetylations are important for transcriptional regulation. The acetylation of lysine residues in histones neutralizes the charge of the side chain and thus changes the ability of histones to bind DNA: this in turn leads to changes in the three-dimensional structure and accessibility of the DNA. While the relative abundance of a particular histone modification may be informative with regard to the general state (gene activation or repression), acetylation dynamics in a specific site may provide detailed insight into the functional activity in its vicinity. The analysis of histone acetylation dynamics using metabolic labelling approaches is thus important to understand this type of epigenetic regulation.

Figure 2a shows a measured spectrum of the fully acetylated histone H3 peptide H3(18-26) from a ^13^C-glucose labelling time course. This peptide can be modified at two lysine residues (K18, K23). After 24 h, approx. 2/3 of the fully acetylated peptide carried an ^13^C-labelled acetyl-group in at least one of the two sites (Table 2) causing a mass shift of 2 Da (M+2, labelling of a single site) or 4 Da (M+4, both sites ^13^C-labelled). In the raw spectrum (Figure 2a) some additional peaks can be observed that are removed after correction for natural isotopes (Figure 2b). Even though some higher mass isotopologues remain, PICor achieves a more accurate correction than AccuCor especially with respect to the lower mass isotopologues. Figure 2c shows the time course measurements for the 3 expected isotopologues (M+0, M+2, M+4) for the fully acetylated H3-peptide at time 0 and 24 hours after label addition, both before and after correction. The relative abundances and calculated time course slopes are in addition listed in Table 2. The isotope correction step removes all isotopologues that are unexpected at time point 0 with 100 % of the abundance stemming from the unlabelled isotopologue (M+0). The effect of isotope correction is highest for the isotopologue with one labelled acetyl-group (M+2). At time point 0 the M+2 value is reduced to 0 %, resulting in a 46 % change of the time course slope from 1.3 % per hour to 1.9 % per hour.

**Table 2:**
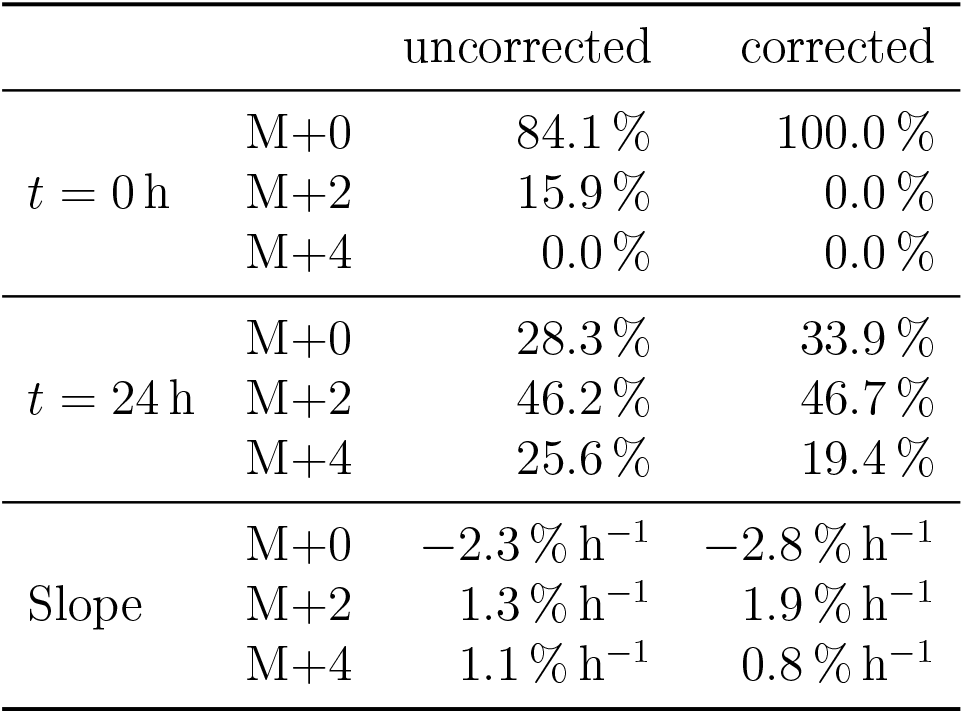
Relative abundances and calculated slopes of uncorrected and corrected acetylation timecourse data of the fully acetylated histone 3 peptide H3(18-26) resulting in 3 different isotopologues with either none (M+0), one (M+2) or two (M+4) acetyl groups labelled.

**Figure 2:**
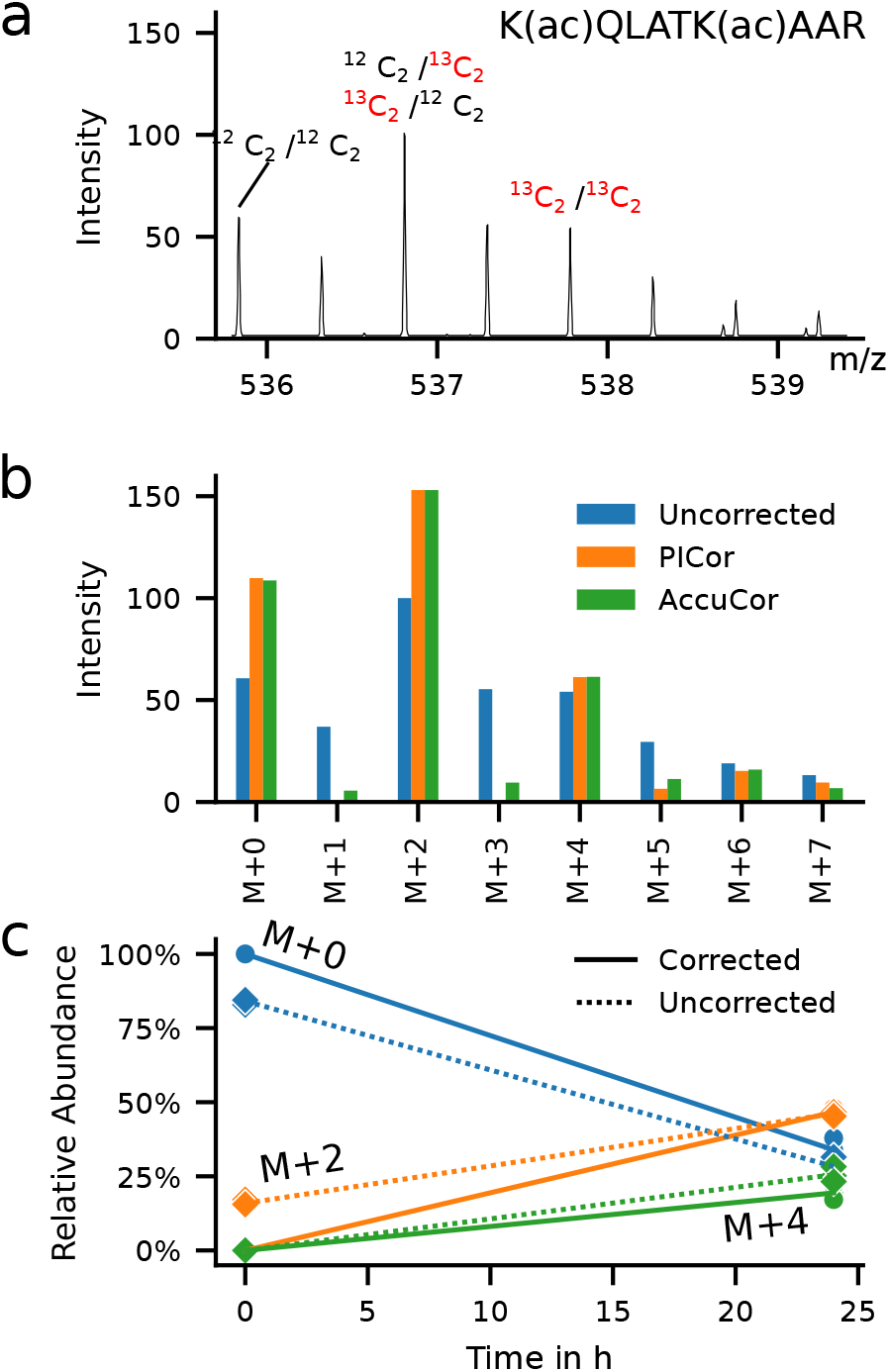
Metabolic labelling of fully acetylated histone 3 peptide H3(18-26) using ^13^C-glucose: a) Example spectrum of time point 24 h.b) Comparison of uncorrected peak height from the spectrum shown in a) and corresponding values corrected with PICor and AccuCor c) Comparison of uncorrected and PICor corrected time course data. There are three different possible labelling states of the fully acetylated peptide. As either none (M+0), one (M+2) or both (M+4) acetyl-groups at the two lysine residues (K18 and K23) in the H3(18-26) peptide can be labelled with ^13^C. Solid line – isotope corrected data; dashed line – uncorrected data; round dots – data point after correction; diamond dots – data point before correction.

### Potential application in quantitative proteomics

Besides metabolic labelling of proteins and protein modifications, stable isotopes are routinely applied in quantitative proteomics to investigate differentially expressed proteins. For example in SILAC (**S**table **I**sotope **L**abeling with **A**mino acids in **C**ell culture) experiments, cells are cultured in media containing either unlabelled, ^13^C labelled or ^13^C and ^15^N labelled lysine or arginine. SILAC is not only used for a quantitative comparison of protein abundance under different experimental conditions, but has also been used to investigate protein turnover^3^. To date it is not common to correct for natural isotopes in SILAC based quantitative proteomics. The expected error is lower compared to metabolic labelling of histone acetylations as the mass difference is larger in most cases and thus the likelihood of naturally labelled peptides with the same mass much lower. Nevertheless, with increasing accuracy and precision of the measurements, appropriate isotope correction will become critical to improve the analysis of these types of experiments.

High-resolution mass spectrometry approaches now enable the analysis of metabolite and protein modification dynamics using metabolic stable isotope labelling techniques. Due to the high number of atoms in peptides, the amount of naturally labelled peptides especially when using ^13^C is relatively large compared to small metabolic compounds. Therefore, isotopologue correction is essential to correctly estimate protein modification dynamics. PICor is a tool developed to enable isotopologue correction for both metabolites and proteins to improve the comprehensive analysis of complex labelling experiments.

## Declarations

### Funding

JD, MZ and IH have been supported by the The Research Council of Norway (ES633272, 302314). MK has received funding from the European Respiratory Society (ERS, RESPIRE3, project no.: R3201703-00121), the Tyrolian Research Fund (project no: 18903) and the Promotion program for Young Scientists at University of Innsbruck (project no: 316826).

### Conflict of interest

The authors declare that they have no conflict of interest.

### Availability of data and material

The data shown is available here:

https://github.com/MolecularBioinformatics/paper_PICor

### Code availability

The PICor source code can be found here: https://github.com/MolecularBioinformatics/PICor

### Human and animal rights

Not relevant.

## Experimental

### Chemicals

Water, acetonitrile (ACN), formic acid (FA, all HPLC-grade) were obtained from Thermo Fisher Scientific (Dreieich, Germany). All other chemicals were purchased from Sigma-Aldrich (Munich, Germany) unless otherwise noted.

### Metabolic labelling with uniformely labelled ^13^C-glucose

The murine macrophage-like cell line RAW264.7 (American Type Culture Collection, Wesel, Germany) or HEK293 cells were cultured in 6-well plates at 37°C and 5% CO2 atmosphere in DMEM with 4.5 g/L glucose. Media was supplemented with 2 mM extra L-Glutamine and 10% (v/v) heat-inactivated FBS. For the labelling experiment, cells were first cultured in media containing uniformly ^12^C-glucose ([U–^12^C]-Glc) for 38 hours. Subsequently, the culture media was replaced by [U–^13^C]-Glc (Euroisotope, Saarbrücken, Germany) containing media and the cells were cultured for up to 24 hours. The cells were harvested by scraping before the exchange of culture medium (t= 0h) or after incubation with [U–^13^C]-Glc containing media for the indicated time.

### Extraction of histone proteins from cells

For histone extraction, the cells were washed twice with a PBS solution (T= 37°C) and lysed by adding 1mL of ice-cold lysis buffer (PBS, 0.5% Triton X-100 (v/v), 1 mM phenylmethylsulfonyl fluoride (PMSF), 1 mM sodium butyrate, 10 *µ*M suberanilohydroxamic acid (SAHA), 10 mM nicotinamide). The cells were subsequently sonificated on ice (3 tunes, 10 sec. 50%) and centrifuged for 10 min at 10,000 rcf at 4°C. After aspirating the supernatant, the pellets were resuspended in 0.5 mL ice-cold resuspension buffer (13 mM EDTA, 10 mM 2-Amino-2-(hydroxymethyl)-1,3-propanediol (Tris base), pH 7.4). After centrifugation (10 min at 10,000 rcf, 4°C) and aspiration of the supernatant, pellets were resuspended in 200 *µ*L ice-cold HPLC-H_2_O. Sulfuric acid (H_2_SO_4_) was added to the samples to a final concentration of 0.4M. Samples were incubated on ice for 1 hour on a shaker using 400 rpm. After centrifugation (10 min at 10,000 rcf, 4°C), the supernatant containing the histone proteins, was added to 1.5 mL acetone and left at -20°C overnight to precipitate the proteins. After centrifugation, acetone was removed and the pellet was dried at room temperature. The pellet was subsequently redissolved in 50 *µ*L HPLC-H_2_O. Protein concentration was determined using the microplate BCA protein assay kit (Thermo Fisher Scientific, Dreieich, Germany) following the manufacturer’s instructions.

### Tryptic in-solution digestion of histone samples

For tryptic digestion, 6 *µ*g of the histone extract were diluted in HPLC-H_2_O to a final volume of 50 *µ*L. For reduction, the samples were incubated with 10 mM dithiothreitol (DTT, dissolved in 100 mM ammonium bicarbonate (ABC), pH 8.3) for 10 min at 57°C on a shaker (600 rpm). Afterwards, the samples were alkylated with 20 mM iodacetamide (IAA, dissolved in 100 mM ABC, pH 8.3) and incubated for 30 min at room temperature in the dark. Subsequently, free IAA was quenched by adding 10 mM of DTT (dissolved in 100 mM ABC, pH 8.3). For tryptic digestion, the samples were incubated with 1 *µ*L of a trypsin solution (c = 0.2 *µ*g/*µ*L sequencing grade modified trypsin, dissolved in trypsin resuspension buffer, Promega, Walldorf, Germany) for 16 hours at 37°C. Afterwards, the samples were acidified by adding formic acid (FA) to final concentration of 0.1% FA.

### LC-MS/MS analysis and preprocessing of histone samples

For LC-MS/MS analysis, 100 ng of the tryptic peptide digests were injected on a nano-ultra pressure liquid chromatography system (Dionex UltiMate 3000 RSLCnano pro flow, Thermo Scientific, Bremen, Germany) coupled via electrospray-ionization (ESI) source to a tribrid orbitrap mass spectrometer (Orbitrap Fusion Lumos, Thermo Scientific, San Jose, CA, USA). The samples were loaded (15 *µ*L/min) on a trapping column (nanoE MZ Sym C18, 5*µ*m, 180umx20mm, Waters, Germany, buffer A: 0.1% FA in HPLC-H_2_O; buffer B: 80% ACN, 0.1% FA in HPLC-H_2_2O) with 5% buffer B. After sample loading, the trapping column was washed for 2 min with 5% buffer B (15 *µ*L/min) and the peptides were eluted (250 nL/min) onto the separation column (nanoE MZ PST CSH, 130 A, C18 1.7*µ*, 75*µ* mx250mm, Waters, Germany) and separated with a gradient of 5–30% B in 25 min, followed by 30-50% in 5 min. The spray was generated from a steel emitter (Fisher Scientific, Dreiech, Germany) at a capillary voltage of 1900 V. For MS/MS measurements, an inclusion list was used for the two times acetylated isotopologue species of the H3 peptide KQLATKAAR. The fragment spectra were acquired in the orbitrap mass analyzer using a resolution of 15,000 (at m/z 200), a normalized HCD collision energy of 30, an AGC target of 5e4 and a maximum injection time of 120 ms. The full MS spectra were acquired in the orbitrap mass analyzer with a resolution of 60,000 (at m/z 200), an AGC target of 4e5, a maximum injection time of 50 ms and over a m/z-range of 350-1650 using the ETD source for internal mass calibration.

Data analysis was performed in FreeStyle 1.6 (Version 1.6.75.20, Thermo Scientific, Bremen, Germany). For the three two-times acetylated KQLATKAAR isotopologues (K(ac12C)QLATK(ac12C)AAR, m/z: 535.8195; K(ac12C)QLATK(ac13C)AAR, m/z: 536.8228; K(ac13C)QLATK(ac13C)AAR, m/z: 537.8262), extracted ion chromatograms (EIC) of all isotopes of the isotopic distributions of the different isotopologues were generated with a mass tolerance of 3 ° ppm. For peak detection, the Genesis algorithm was used with the following parameters: percent of highest peak: 10, minimum peak height (signal/noise): 5, signal-to-noise threshold: 3, tailing factor: 3. The peak area and the peak height was exported for each isotope of the isotopic distributions of the three different isotopologues to a tab-delimited text file.

### Acetyl Co-A extraction

For metabolite extraction, HEK293T cells were washed three times with ice-cold PBS solution. 300 *µ*L of ice-cold methanol were added to the cells to stop metabolic reactions followed by 300 *µ*L of water. Cells were scraped and transferred to an Eppendorf tube containing 300 *µ*L of chloroform. Afterwards, the samples were incubated for 20 min on a thermomixer (1400 rpm, 4°C). Phase separation was achieved by centrifugation (5 min, 16000 rcf, 4°C). 450 *µ*L of the methanol phase were used to extract acetyl-CoA by solid phase extraction (SPE) using a (2-(2-Pyridyl)ethyl silica gel based SPE column (Supelco, Merck, Sigma Aldrich, Germany) and a vacuum extraction manifold (Waters 20-Position Extraction Manifold, Waters, Austria). For SPE extraction, the columns were equilibrated with 1 mL of equilibration buffer (45% ACN, 20% H2O, 20% Acetic Acid, 15% Isopropanol (v/v), pH 3). After equilibration, samples (polar methanol phase) were loaded on the SPE column, followed by a washing step using 1 mL of the equilibration buffer. Analytes were eluted from the SPE columns with 2 mL of MeOH/250 mM NH4FA (4+1 v/v, pH 7). The eluates were dried and stored at -80°C for further LC-MS analysis.

### Measurement and analysis of acetyl-CoA

Dried samples were re-dissolved in 100 *µ*L of 50% ACN. For LC-MS/MS analysis, 1 *µ*L of sample were injected on a ultra pressure liquid chromatography system (Vanquish Flex Quarternary UHPLC System, Thermo Scientific, Bremen, Germany) with a hydrophilic interaction liquid chromatography (HILIC) column (Aquity UPLC BEH Amide, 130Å, 1.7*µ*m, 2.1 × 150mm, Waters, Germany). The UPLC was coupled via an electrospray-ionization (ESI) source to a QExactive HF Orbitrap (Thermo Scientific, Bremen, Germany). Chromatographic separation was performed with a gradient going from 95%-50% B in 8 min, and then from 50%-10% B in 2 min (A: 10 mM NH4Ac in H2O, pH 10; B: 95% ACN, 5% 10 mM NH4Ac in H2O, pH 10). MS analysis was carried out with targeted single ion monitoring (SIM) scans in positive mode. SIM windows were set to include all possible isotopologue forms of Acetyl-CoA up to M+23. The SIM window for Acetyl CoA was set at a retention time between 9.2 min and 11 min and the m/z isolation window was set to 821.733 ° 15. A resolution of 60,000 FWHM at 200 m/z was used. The maximum injection was set to 80 ms and the AGC target to 5e4.

Data analysis was performed in TraceFinder 5.0 (Version 5.0.889.0, Thermo Scientific, Bremen, Germany). Peaks were fitted using the Genesis algorithm with the following parameters: percent of highest peak: 1, minimum peak height (signal/noise): 3, signal-to-noise threshold: 2, tailing factor: 1. Peak integration was manually corrected if necessary.

### Natural isotope correction

We used the skewed matrix approach for correction of high resolution natural isotope abundance as described by Heinrich et al. 2018^12^. The resolution correction takes all isotopologues into account that are within the resolving limits at a given resolution according to Su et al. 2017^7^. PICor is a Python (version 3.8.5) based implementation of the isotopologue correction tool and is freely available on GitHub https://github.com/MolecularBioinformatics/PICor and can be used for any organic compound. The user needs to specify the chemical composition and the type of labelling experiment and supply a data file with the peak integration values from MS measurements for the isotopologues of interest. It is not required to provide measurements for all possible isotopologues, which makes it applicable for proteomics data. Full documentation can be found in the GitHub repository.

### Histone acetylation analysis

To get the inital slope of the acetylation reactions in Table 2, the SciPy (version 1.5.2)^14^ *linregress* function was used to fit the relative abundance of the respective substrates with respect to time to linear equations.

The data was corrected with PICor using the *resolution_correction* option with a resolution of 60,000 at m/z 200 (*mz-calibration* parameter).

### Software for isotopologue correction

The following software versions were used for the comparison of isotopologue corrections:

- AccuCor: v0.2.3.9000
- IsoCor: v2.2.0
- IsoCorrectoR: v1.6.2
- PICor (our tool): v0.6.2

## Notes

### Competing Interest Statement

The authors have declared no competing interest.

### Summary of Updates

This is a considerable revised version of the manuscript including new experimental data and an update of the algorithm underlying the isotopologue correction tool PICor.

